# Impact of cardiometabolic factors and blood beta-amyloid on the volume of white matter hyperintensities in dementia-free subjects with cognitive complaints

**DOI:** 10.1101/2024.06.10.598296

**Authors:** Patricio Riquelme-Contreras, Michael Hornberger, Mizanur Khondoker, Fernando Henriquez, Cecilia Gonzalez-Campo, Florencia Altschuler, Matías Fraile-Vazquez, Pamela Lukasewicz-Ferreira, Bruna Bellaver, Thomas Karikari, Tharick A. Pascoal, Graciela Muniz-Terrera, Cecilia Okuma, Christian Gonzalez-Billault, Felipe Court, Mauricio Cerda, Patricia Lillo, Andrea Slachevsky

**Affiliations:** Geroscience Center for Mental Health and Metabolism (GERO Chile), Chile; Department of Medical Technology. Faculty of Medicine. Universidad de Chile, Chile; Laboratory of Neuropsychology and Clinical Neurosciences (LANNEC). ICBM. Faculty of Medicine, Universidad de Chile, Chile; Masters in Biological Sciences / Neuroscience program. Faculty of Sciences. Universidad de Valparaíso, Chile; Norwich Medical School, University of East Anglia, Norwich, UK; Laboratory for Cognitive and Evolutionary Neuroscience (LaNCE), Department of Psychiatry, Faculty of Medicine, Pontificia Universidad Católica de Chile, Chile; Cognitive Neuroscience Center, Universidad de San Andrés, Argentina; National Scientific and Technical Research Council (CONICET), Buenos Aires, Argentina; Department of Psychiatry, University of Pittsburgh, Pittsburgh, PA 15213, USA; Ohio University Heritage College of Osteopathic Medicine, Athens, OH, USA; Edinburgh Dementia Prevention, Centre for Clinical Brain Sciences, University of Edinburgh, Edinburgh, UK; Center for Medical Informatics and Telemedicine (CIMT), Faculty of Medicina, Universidad de Chile, Chile

**Keywords:** white matter hyperintensities, dementia-free cognitive complaint individuals, Aβ blood biomarkers, cardiometabolic factors

## Abstract

**Introduction:** White matter hyperintensities (WMH) in Alzheimer’s disease (AD) have traditionally been associated with cerebrovascular diseases. Amyloid β (Aβ) deposition reportedly contributes to WMHs; however, this relationship remains unclear in dementia-free subjects with cognitive complaints (CC). Here, we explored the relationship between WMHs and cardiometabolic and Aβ blood biomarkers in a community-based cohort of Latin American CC participants.

**Methods:** We recruited 112 individuals with CC (69 – 92 YO, 90 females) with available plasma Aβ biomarkers and cardiometabolic markers (systolic – diastolic blood pressure and glycaemia). WMHs were quantified using a lesion segmentation tool based on SPM12 and segmented using the John Hopkins University (JHU) Atlas and ALVIN segmentation for periventricular and subcortical white matter. Linear multiple regression models were fitted to assess total WMH lesions and the segmented tract, using demographics, cardiometabolic, and Aβ blood biomarker measures as independent variables.

**Results:** After multiple comparison corrections, diastolic blood pressure was associated with WMHs, specifically in the right anterior thalamic radiation, left cingulum, minor forceps, and subcortical ALVIN segmentation. Glycaemia was associated with WMH volume in forceps major, forceps minor, and right fronto-occipital fasciculi. Conversely, Aβ blood biomarkers and systolic blood pressure showed no association with WMH overall or in specific tracts.

**Conclusion:** Our findings suggest that, in dementia-free CC individuals, WMH volume was more related to cardiometabolic factors, whereas Aβ blood biomarkers might be of less relevance. Dementia prevention strategies in individuals might be a useful focus for managing high peripheral vessel resistance and endothelial damage due to hypertension and hyperglycaemia.

## Introduction

Dementia is highly prevalent in the aging population, with Alzheimer’s disease (AD) being the most common form [1]. AD progresses along a continuum, from cognitive complaints (CC) to dementia. There is now substantial evidence to indicate that earlier detection of AD provides an opportunity to slow disease progression through the administration of medication and implementation of lifestyle changes. However, there is currently no cure for AD, and most therapies are focused on providing only modest relief for symptomatic stages of the disease. Consequently, there is a significant motivation to identify the disease in its preclinical stages [2] as studying prevention and therapy at these stages may provide better opportunities for therapeutic success.

Currently, the distinction between normal aging and CC and its progression to AD is subtle [3]. There is ongoing work on dementia biomarkers to detect the earliest changes associated with AD before clinical symptoms can be detected via clinical evaluation and neuropsychological tests to establish whether these symptoms represent prodromic stages of the disease, particularly if CC non-demented individuals are in the early stages of AD dementia. Individuals without dementia with CC can be broadly classified as having subjective cognitive decline (SCD) or mild cognitive impairment (MCI), based on neuropsychological tests or informant-based surveys [4], [5], [6]. The rationale for SCD is that, in the absence of objective neuropsychological deficits, individuals can perceive a change in their memory and/or other cognitive abilities relative to their previous level of performance [7], [8], while MCI is characterised by a CC associated with an objective cognitive impairment. Nevertheless, the natural history and disease mechanisms of AD and related disorders remain poorly understood. Scarce resources are available to scrutinise patients as early as needed and the use of integrative approaches combining standardised, repeated clinical investigations and cutting-edge biomarker measurements is limited [9].

There are currently several ongoing investigations into how neuroimaging biomarker signatures may corroborate CC as part of the AD continuum [4]. This research has identified useful biomarkers for identifying structural brain changes related to CC. White matter hyperintensities (WMHs) are particularly important in this context given their high prevalence in the aging population, as well as their potential role as early indicators of dementia pathophysiology, although their pathogenesis is not well understood.

WMH are commonly detected using Fluid-Attenuated Inversion Recovery (FLAIR) magnetic resonance imaging (MRI). In FLAIR images, areas of higher signal intensity indicate prolonged relaxation times owing to an increase in bound water within the tissue [10]. WMH have further been associated with an increased risk of general dementia and AD in the general population [11], [12], although their exact contribution to the pathophysiology of dementia is still being explored (Murray et al., 2012). WMH are considered a hallmark feature of subclinical cerebrovascular disease [13], that may eventually lead to long-term neurological deficits or cognitive decline [14]. In this context, traditional risk factors for cardiovascular diseases (hypertension, diabetes, and others), including lifestyle behaviours such as smoking, diet, and physical activity, have been linked to the emergence of WMHs [14], [15]. For example, hypertense individuals often have significantly worse cognitive function, which is associated with WMH load (García□Alberca et al., 2020). Similarly, diabetes mellitus has been associated with a greater overall WMH burden [16]. Hypertension, diabetes, and other lifestyle factors are known to damage cerebral vessels through multiple mechanisms (mostly affecting blood-brain barrier integrity and neuroinflammatory processes), ultimately resulting in the emergence of WMHs, along with other cerebrovascular pathologies (e.g. lacunes, microbleeds, and enlarged perivascular space)[16], [17]. However, cardiometabolic factors may not be the only contributors to the development of WMH. There have also been suggestions that Aβ and tau accumulation in preclinical AD may also contribute to the occurrence of WMHs [18]. Indeed, recent findings have suggested that the primary origin of WMHs is due to Aβ and tau pathology and should be considered as main contributors to WMHs [19], [20]. Specifically, WMHs are more commonly associated with Aβ PET deposition than with tau PET burden (Graff-Radford et al., 2019). Indeed, some reports have indicated a positive association between Aβ PET deposition and WMH in the middle temporal and fusiform regions[21].

Furthermore, there is evidence to suggest that while WMHs are primarily associated with cardiometabolic risk and neurodegeneration, AD-specific pathways may contribute to their formation in a regionally specific manner. For example, phosphorylated CSF tau (p-tau) has been linked with temporal lobe WMHs, while CSF Aβ (Aβ 42/Aβ 40 ratio) has been associated with parietal lobe WMHs[22]. These findings raise the question of whether the observed WMHs are due to cardiometabolic risk factors, AD pathophysiology, or a mixture of both in preclinical AD, such as CC.

The current study aims to address this shortcoming by investigating the relationship between WMHs and: i) AD blood biomarkers (Aβ), as well as ii) cardiometabolic risk factors (arterial blood pressure and glycaemia) in a community-dwelling cohort with non-demented CC. We further hypothesised that cardiometabolic factors would largely determine the WMH load in SCD with AD blood biomarkers, with only a small additional effect on the WMH load.

## Material and methods

### Participants

We included 112 individuals (90 females) selected from a Chilean CC non-demented community-based cohort (GERO Chile)[23]. The inclusion criteria for this cohort were adults > 70 years old, not diagnosed with dementia, with self-declared cognitive complaints or those declared by an informant, and home-dwelling (not living within a care facility). Participants were identified through a household census, and the evaluation protocol was based on a multidimensional approach, including sociodemographic, biomedical, psychosocial, neuropsychological, neuropsychiatric, and motor assessments. Neuroimaging, blood, and stool samples were also obtained (see [23] for an exhaustive description of the protocol). All participants underwent neurological evaluation to verify that they fulfilled the inclusion criteria.

The neuropsychological evaluation in this cohort included the Montreal Cognitive Assessment (MoCA) test, validated for the Chilean population, for which the optimal cut-off point for MCI was ≤ 20, with sensitivity and specificity rates of 75% and 82% for aMCI and 90% and 86% for mild dementia, respectively [24]. In addition, all of these individuals had the following available data: total WMH lesions in FLAIR MRI images, as well as the following systemic biomarkers available in the database: systolic blood pressure, diastolic blood pressure, glycaemia, and the blood biomarkers Aβ42 and Aβ40. Based on Aβ indexes, we calculated the Aβ42/40 ratio, which was used for the final analysis.

The GEROChile project was approved by the Scientific Ethics Committee of the Eastern Metropolitan Health Service of Santiago de Chile. Informed consent was obtained from all the participants, as approved by the same committee.

### MRI acquisition and processing

MRI T1W and FLAIR MR images were obtained with the following acquisition parameters: T1wMPRAGE: TR: 1710 ms, TE: 2.25, FoV: 224 mm, voxel size: 1.0 × 1.0 × 1.0, slice thickness: 1.0 mm, flip angle: 8°, and FLAIR: TR: 8000 ms, TE: 90 ms, TI: 2500 ms, FoV: 220 mm, voxel size: 0,7 x 0,7 x 4,0 mm, slice thickness: 4,0 mm, flip angle: 150°. Images were acquired using a Skyra 3T scanner (Siemens) at the Neurosurgery Institute of Dr. Alfonso Asenjo (Santiago, Chile).

The images were preprocessed for movement correction and field inhomogeneities and registered in the subject’s space using structural T1w images. WMH segmentation was performed using the lesion prediction algorithm (LPA) (http://www.applied-statistics.de/lst.html) implemented in the LST toolbox version 2.0.15, for SPM12 (https://www.fil.ion.ucl.ac.uk/spm). To obtain better segmentation results, individual FLAIR images were used as the only input to obtain lesion probability maps[25]. These maps were visually inspected per subject to check for possible artefacts and were discarded when artefacts were found. The probability maps were thresholded using a default value of 0.1 and normalized. The total WMH volume (mm^3^) was extracted from the subject-level WMH maps. Subsequently, WMH were segmented based on two white matter fibre bundle atlases: the John Hopkins University white matter tractography atlas [26], [27], [28], for which 20 tracts were identified probabilistically by averaging the results of running deterministic tractography on 28 normal subjects; and the ALVIN (Automatic Lateral Ventricle Atlas DelIneatioN) atlas, a fully automated algorithm which works within the SPM to segment lateral ventricles from structural MRI images, and has been validated in infants, adults, and patients with Alzheimer’s disease (ICC > 0.95). ALVIN is insensitive to different scanner sequences (ICC > 0.99, 8 different sequences 1.5T and 3T), but sensitive to changes in ventricular volume [29]. This segmentation identified periventricular and subcortical white matter regions. Finally, all measurements were normalised to the total intracranial volume (TIV) for comparison. This procedure was performed automatically using the VBM8 Toolbox of SPM12, which segments the voxels of T1-weighted images into grey matter, white matter, and cerebrospinal fluid. The sum of these values represents the TIV. As an outcome of this process, we obtained the total lesion load of the WMH, as well as the lesion load of the WMH per segmented tract (in mm3), based on the atlases described and normalised by the total intracranial volume.

### Blood samples and cardiovascular measurements

Systolic and diastolic blood pressure were measured during the cardiovascular cohort evaluation. These indices were obtained at rest from one measurement on the left arm without crossing the legs, with the feet completely supported on the floor. Both arms were supported by a closed surface. Measurements were performed using a digital blood pressure monitor model UA-611A (A&D Instruments Limited, UK). Glycaemia samples while fasting were obtained in the subject’s home through a vacuum system with venous access of 21 G or 23 G, depending on the subject’s venous calibre. These samples were processed at the Central Laboratory of the Hospital del Salvador, Santiago, Chile.

For neurodegenerative biomarker assessment, blood was collected in a separate session, one week after glycaemia acquisition and after an overnight fast. Samples were divided into aliquots and frozen at -80°C. These samples were the sent to Pittsburgh, Pennsylvania, to conserve the cold chain and under permanent temperature monitoring until processing. Plasma biomarkers were quantified using a Neurology 4-Plex E (#103670) commercial assay kit (Quanterix, Billerica, MA, USA).

### Statistical analysis

Demographic and clinical descriptive statistics were used to estimate the mean, standard deviation (SD), min, and max. The variables included sex, age (years), education (years), available biomarkers, and WMH lesion load. To establish the relationship between WMH lesion load and selected biomarkers, we used a linear multiple regression model to fit the total lesion load WMH per segmented tract. Post-hoc Holm– Bonferroni test was performed for multiple comparison correction. For a better representation of the model, the lesion load of the WMH was transformed from mm^3^ to μm^3^ and then, to a natural logarithmic scale. The variables used to fit the model were age (years), sex, years of formal education, Aβ42/40 ratio, systolic and diastolic blood pressure, and glycaemia.

We applied traditional threshold levels of p < 0.05 for the total model and per specific biomarker. All statistical procedures were performed using the STATA/SE 17.0, software package (StataCorp LLC, https://www.stata.com).

## Results

### Demographics, lifestyle factors and AD biomarkers

The average age of the participants was 76.32 years (SD = 5.18), with an average years of formal education of 9.28 years (SD = 4.72). The average MoCA test score was 21.64 (SD = 4.47). The cutoff score for the Chilean adaptation of this neuropsychological assessment for SCD to be considered was ≥ 21. These findings confirm that our cohort predominantly aligns with the cognitive complaint spectrum, including SCD and MCI, in individuals without dementia.

Regarding blood pressure and blood glucose levels, our samples exhibited, on average, level 1 systolic hypertension (139.61 mmHg) and normal diastolic blood pressure (73.45 mmHg), according to the American Heart Association Council (https://heart.org/bplevels). The fasting glycemia average was 96.29 mg/dL, which is considered normal, considering that the threshold to determine hyperglycaemia according to the CDC guidelines is 100 mg/dL (https://www.cdc.gov/diabetes/basics/getting-tested.html).

### Amyloid - beta biomarkers

The average blood concentration of Aβ42 was 6.643 pg/mL (SD = 1.808), while that for Aβ40 was 109.39 (SD = 21.476). Even though reference intervals for plasma biomarkers of AD have not yet been established, some evidence has established that the reference intervals for Aβ42 and Aβ40 are between 2.72–11.09 pg/mL and 61.4–303.9 pg/mL, respectively. The average for Aβ42/40 was 0.059 (SD = 0.012), and the reported reference interval for this index was 0.022–0.064[30]. Based on this evidence, our group presented normal levels of these biomarkers.

The demographic, cardiometabolic and Aβ biomarkers summary statistics are presented in Table 1.

**Table 1:**
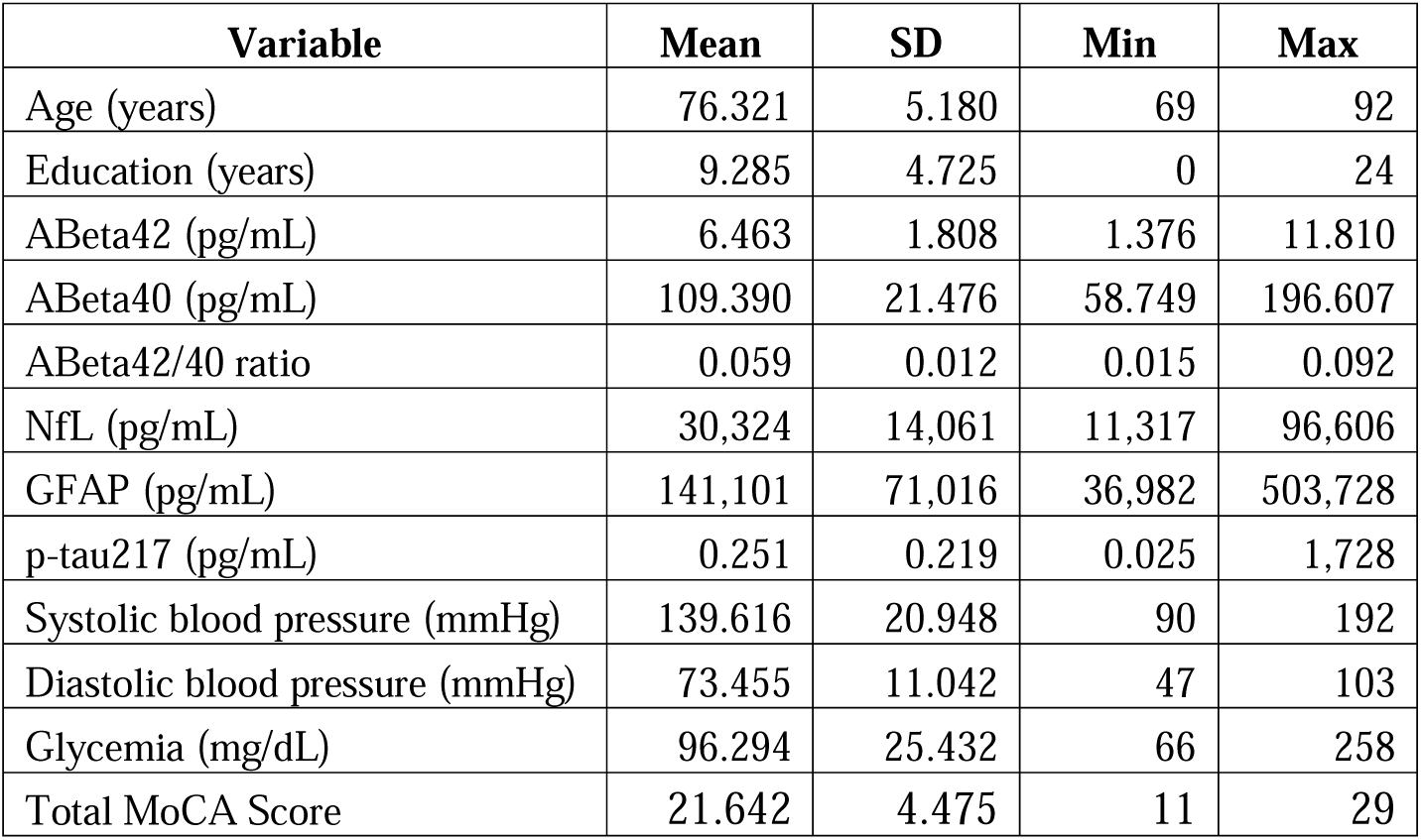
Demographics and clinical characteristics of the study participants (n = 112).

### White Matter Hyperintensities

The volume of WMHs was highly variable across all tracts, with the thalamic radiations (left: mean = 1287324 μm^3^, SD = 908302; right: mean = 868533 μm^3^, SD = 555540) and corpus callosum (forceps major: mean = 1055505 μm^3^, SD = 780131; forceps minor: mean = 532523 μm^3^, SD = 416197) having the highest WMH lesion volumes, while left cingulum hippocampal (mean = 8938.3 μm^3^, SD = 18185) and right temporal superior longitudinal fasciculus (mean = 7883 μm^3^, SD = 23588) showed the lowest mean WMH lesion load.

For the ALVIN segmentation, the periventricular region showed a higher WMH lesion volume (mean = 5898777 μm3, SD = 5278707) compared to the subcortical regions (mean = 3901206 μm^3^, SD = 5074165). Descriptive statistics for all segmented tracts in both atlases are presented in Tables 2 and 3.

**Table 2:**
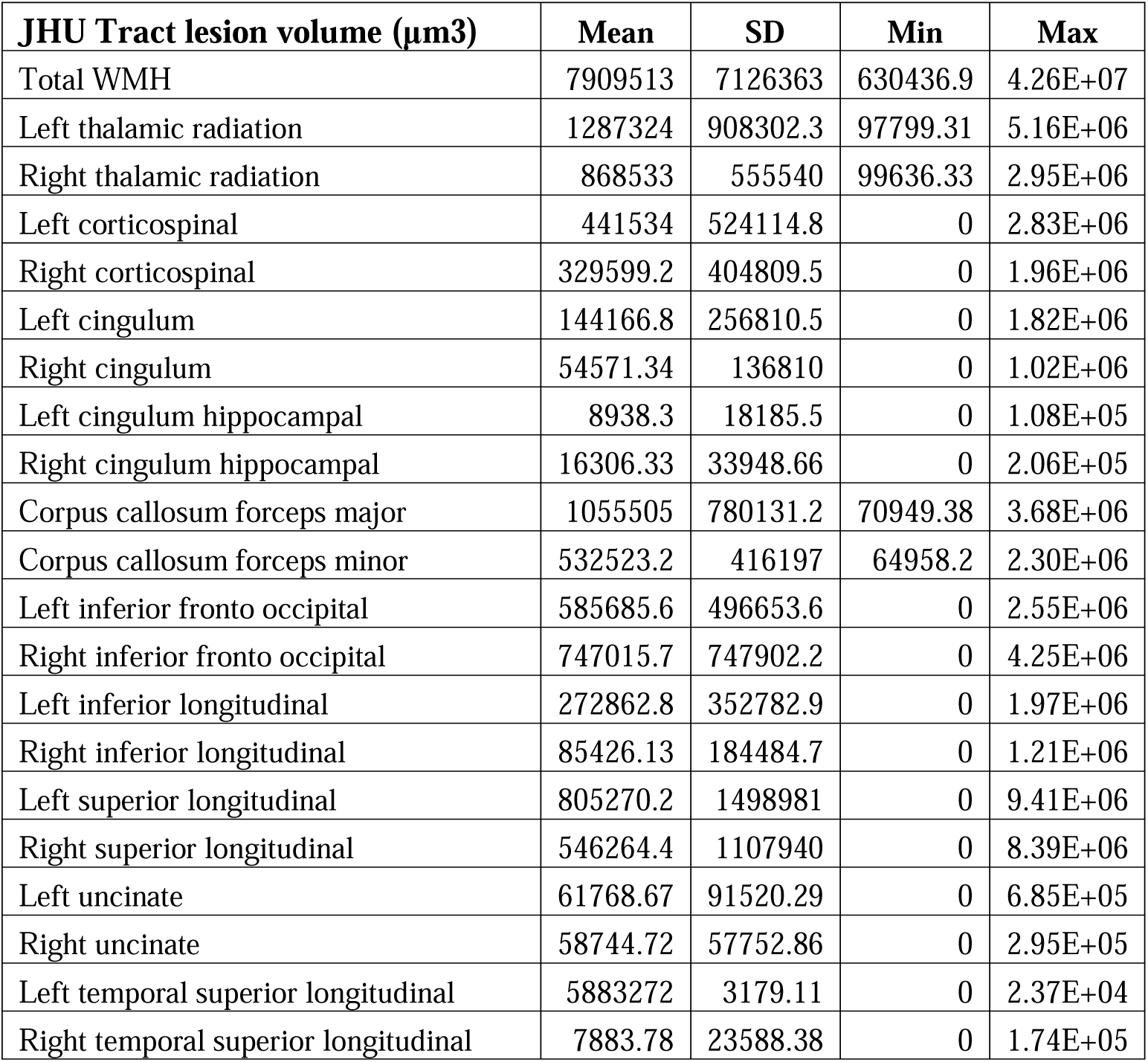
Descriptive statistics for all tracts based on John Hopkins University (JHU) white matter segmentation.

| **JHU Tract lesion volume ($\mu\text{m}^3$)** | **Mean** | **SD** | **Min** | **Max** |
| --- | --- | --- | --- | --- |
| Total WMH | 7909513 | 7126363 | 630436.9 | 4.26E+07 |
| Left thalamic radiation | 1287324 | 908302.3 | 97799.31 | 5.16E+06 |
| Right thalamic radiation | 868533 | 555540 | 99636.33 | 2.95E+06 |
| Left corticospinal | 441534 | 524114.8 | 0 | 2.83E+06 |
| Right corticospinal | 329599.2 | 404809.5 | 0 | 1.96E+06 |
| Left cingulum | 144166.8 | 256810.5 | 0 | 1.82E+06 |
| Right cingulum | 54571.34 | 136810 | 0 | 1.02E+06 |
| Left cingulum hippocampal | 8938.3 | 18185.5 | 0 | 1.08E+05 |
| Right cingulum hippocampal | 16306.33 | 33948.66 | 0 | 2.06E+05 |
| Corpus callosum forceps major | 1055505 | 780131.2 | 70949.38 | 3.68E+06 |
| Corpus callosum forceps minor | 532523.2 | 416197 | 64958.2 | 2.30E+06 |
| Left inferior fronto occipital | 585685.6 | 496653.6 | 0 | 2.55E+06 |
| Right inferior fronto occipital | 747015.7 | 747902.2 | 0 | 4.25E+06 |
| Left inferior longitudinal | 272862.8 | 352782.9 | 0 | 1.97E+06 |
| Right inferior longitudinal | 85426.13 | 184484.7 | 0 | 1.21E+06 |
| Left superior longitudinal | 805270.2 | 1498981 | 0 | 9.41E+06 |
| Right superior longitudinal | 546264.4 | 1107940 | 0 | 8.39E+06 |
| Left uncinate | 61768.67 | 91520.29 | 0 | 6.85E+05 |
| Right uncinate | 58744.72 | 57752.86 | 0 | 2.95E+05 |
| Left temporal superior longitudinal | 5883272 | 3179.11 | 0 | 2.37E+04 |
| Right temporal superior longitudinal | 7883.78 | 23588.38 | 0 | 1.74E+05 |

**Table 3:** Descriptive statistics for all tracts based on the Automatic Lateral Ventricle Atlas DelIneatioN (ALVIN) atlas, a fully automated algorithm which works within SPM to segment the lateral ventricles from structural MRI images.

| **ALVIN segmentation volume (µm<sup>3</sup>)** | **Mean** | **SD** | **Min** | **Max** |
| --- | --- | --- | --- | --- |
| Periventricular | 5898777 | 5278707 | 116813.1 | 3.70E+07 |
| Subcortical | 3901206 | 5074165 | 0 | 3.49E+07 |

This segmentation includes 20 tracts that were identified probabilistically by averaging the results of running deterministic tractography on 28 normal subjects. All measurements are shown in cubic micrometre (μ3). After, measurements were transformed in a natural logarithmic scale to run multiple regression models.

All measurements are shown in cubic micrometres (μm3). Subsequently, measurements were transformed in a natural logarithmic scale to run multiple regression models.

### Relationship of WMH to age, cardiometabolic and amyloid blood biomarkers

We fitted multiple linear regression models to estimate the association of age, cardiometabolic and Aβ blood biomarkers on WMH lesion loads. After Holm-Bonferroni’s multiple comparison correction, age remained the most important factor in explaining overall WMH. We also found that Diastolic blood pressure had a significant association with WMH in right anterior thalamic radiation (β=0.305, p=0.008), right corticospinal tract (β=0.285, p=0.027), left cingulum (β=0.364, p=0.013), corpus callosum forceps minor (β=0.287, p=0.013), and subcortical regions (β=0.328, p=0.007).

Further, glycaemia was found to have a significant association with WMHs in the corpus callosum forceps major (β=0.287, p=0.018), forceps minor (β=0.253, p=0.028), and right inferior fronto occipital fasciculus (β=0.286, p=0.024). However, because most participants were not hyperglycaemic, we conducted a post-hoc analysis only on participants with glycaemia levels of > 100 mg/dL. A total of 35 subjects were included in the post-hoc analysis. In this smaller group, the association did not survive multiple comparisons, likely due to the small sample size. However, there was still a statistical trend for two of the three tracts included in this analysis, even after excluding people without hyperglycaemia, specifically the forceps major (p=0.082) and the inferior fronto-occipital fasciculus (p=0.079).

Notably, neither systolic pressure nor the Aβ42/40 ratio was significantly associated with any WMHs. The regression indices for each fitted model and their adjusted p-values after the post-hoc test are presented in Online Resources 1 and 2. Graphical representations of the results are shown in Figures 1 and 2.

**Fig. 1:**
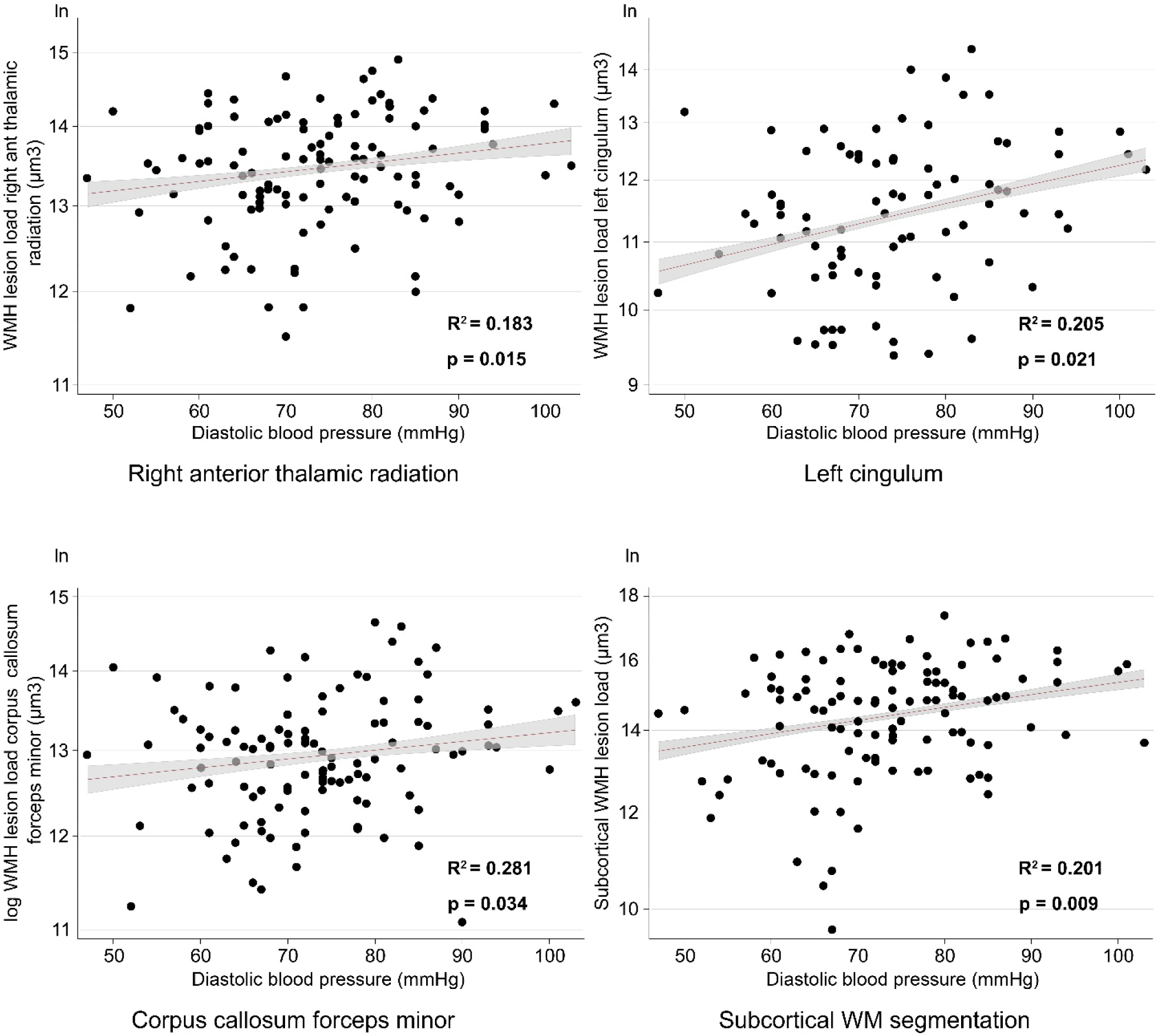
Significant regression models graphics for WMH lesion volume in the function of diastolic blood pressure

**Fig. 2:**
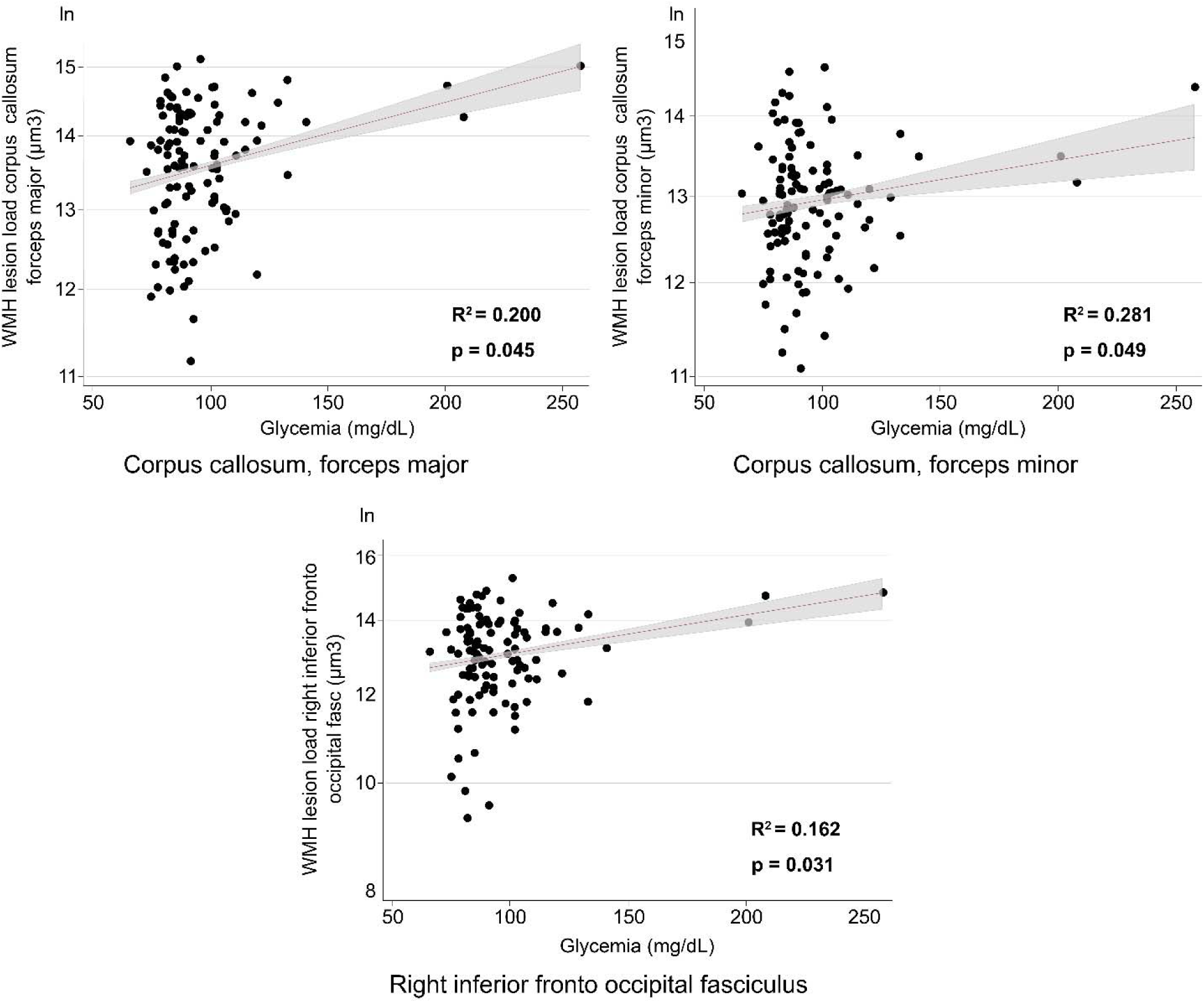
Significant regression model graphics for WMH lesion volume in the function of glycemia

## Discussion

Our study investigated the relationships between WMHs, AD blood biomarkers, and cardiometabolic factors in dementia-free individuals with CC. Our findings suggest that, in addition to age, diastolic blood pressure and glycaemia are associated with WMH lesion load in several white matter fibre tracts. Importantly, plasma Aβ levels were not associated with WMH loads in any tracts. These findings suggest that the WMH load is predominantly driven by cardiometabolic risk factors, and not by incipient amyloid pathophysiology in non-demented subjects with CC.

Hypertension and WMHs are associated with cognitive impairment [31]. WMHs are related to cerebrovascular disorders (Balestrieri et al., 2021), because risk conditions such as hypertension can compromise the cerebral microcirculation via microvascular injury, increased vascular stiffness, increased myogenic tone, microbleeds, blood-brain barrier disruption with neuroinflammation, and glymphatic system impairment [32]. This damage to the microvasculature structure has been associated with cognitive performance and WMH in older hypertensive individuals [33], and is further known to be related to cognitive impairment [34], [35], [36], [37]. Interestingly, in our study, only diastolic blood pressure was associated with WMH lesion volume. The reason for this might be that, although systolic blood pressure linearly increases with age, diastolic blood pressure usually decreases after the age of 55 years, with a shift from mixed or diastolic hypertension to an increased frequency of isolated systolic hypertension in older age [38]. This might explain our results, as persistently high diastolic blood pressure in late life may be more predictive of WMHs than a decrease in systolic blood pressure. However, the exact factors that might explain this dissociation remain unclear and require further investigation.

Our glycaemia findings are also intriguing in this context. It has been well established that hyperglycaemia and type 2 diabetes are associated with white matter lesions (WMLs)[39], increasing WMHs loads [40], [41], [42] and overall white matter microintegrity [40], [43], [44]. The most likely explanation for these hyperglycaemic changes in WMH volume is that they are related to diabetic microangiopathy in small cerebral vessels [45]. Indeed, the pathology of diabetic complications has high similarity with vascular changes, resulting in endothelial dysfunction and atherosclerosis. Diabetes is also a risk factor for vascular diseases and various comorbidities, resulting in the diagnosis of “panvascular disease”[46]. Therefore, our findings regarding glycaemic indices and WMHs are likely due to microvascular compromise, ultimately sharing mechanisms with the effects of diastolic blood pressure on the cerebral microvasculature and vascular peripheral resistance.

In contrast, the results from plasma Aβ biomarkers in WMH remain inconclusive, and the main evidence that links Aβ deposition and WMHs is based on PET, not plasma. One systematic review suggested that PET Aβ accumulation and WMHs are independent, but additive processes [47], whereas other studies have found associations between WMH burden and Aβ biomarkers in cognitively unimpaired adults, MCI, and AD subjects, with Aβ burden in FDG-PET in temporal lobe regions [48], [49]. Most of the evidence related to Aβ plasma biomarkers was based on oligomerized amyloid-β (OAβ) levels and their relationship with WMHs lesion load. WMHs was shown to increase with age in that study, while OAβ levels did not. Further, log-WMHs volume was positively correlated with OAβ (r = 0.24, p = 0.02), and this association was significant in the periventricular area [50].

To our knowledge, our cohort is the first to investigate whether WMHs in CC non-demented people are related to plasma Aβ42/40 biomarkers. According to our results, AD blood biomarkers (represented in this study by the Aβ42/40 ratio) are not associated with WMH volume in CC non-demented individuals. This result is interesting, considering that we analysed data from a South American cohort of CC subjects with sociodemographic features distinct from those of other previously reported cohorts. Instead, we propose that microvascular damage generated by hyperglycaemia and hypertension likely contributes to the emergence of WMHs, such as CC, in the early stages of dementia. We speculate that, based on previous molecular imaging evidence, the effects of this microvascular damage in the brain, such as BBB breakdown, oxidative stress, neuroinflammation, and glymphatic system impairment, could trigger Aβ accumulation and, consequently, tau aggregation in the brain parenchyma, due to the occurrence of microvascular damage when the disease further develops. This hypothesis would be supported by evidence that associates plasma Aβ42 with the presence of cerebral small vessel disease and more advanced cognitive impairments (Qu et al., 2023), although it needs to be further investigated.

In terms of clinical implications, our findings reinforce three main points: 1) Hypertension and glycaemia are related to structural changes in the CC. Thus, the early diagnosis of cardiometabolic risk factors is important in CC, not in demented subjects, to potentially alleviate and influence the progression to MCI, or even dementia. Careful monitoring and management of blood pressure in the elderly and in patients with CC are essential to reduce the incidence and progression of cerebrovascular disease and its consequent cognitive decline [52].

At the same time, management of hyperglycaemic states in old age, as well as the main drivers of diabetes-related cerebral microvascular dysfunction, such as obesity and insulin resistance, are important. Observational studies have suggested that diabetes-related microvascular dysfunction is associated with higher risks of stroke, cognitive dysfunction, and depression. Cerebral outcomes in diabetes might be improved following treatments targeting the pathways through which diabetes damages the microcirculation [53]. Indeed, recent evidence suggests that selective inhibitors of the cytoplasmic enzyme phosphodiesterase-5 (PDE5i), such as sildenafil, vardenafil, and tadalafil, which are vasodilator medications, improve cognitive outcomes before chronic use [54], [55], reinforcing the idea that an increase in cerebral peripheral resistance in the context of microvascular damage could play a role in age-related cognitive decline. Further, MRI FLAIR and susceptibility-weighted imaging (SWI) sequences are useful for detecting changes in white matter hyperintensities in patients with prodromal dementia [56]. Using automated processing tools to quantify microbleeds and WMHs, measurement of Virchow-Robin space, and/or including a standardised visual Fazekas scale for measuring WMHs could ensure the utmost benefit is achieved from this technique in the context of SCD, to allow for the mitigation of potential progression to MCI (Furtner & Prayer, 2021). Further, there is a need for a more integral approach to cardiometabolic factors in preclinical forms of dementia. Elevated blood pressure and hyperglycaemia frequently coexist and are components of metabolic syndrome [57], which, in turn, is related to excessive cortisol secretion due to psychosocial stress-induced hypothalamic-pituitary-adrenal axis activation and Cushing’s syndrome [58], [59]. We propose that a deeper analysis of the effects of glucocorticoids on the cerebral microvasculature and BBB integrity, neuroinflammation, and glymphatic function in both animal and human models could facilitate a more integrated approach to this phenomenon.

### Limitations

Our study is not without limitations. Considering the combination of biomarkers, we used a reasonably large sample for this study. However, it will be important to corroborate our findings using an independent larger sample size. This validation could further improve the accuracy of the fitted models, particularly for patients with higher glycaemic indices. Dysglycaemia was only present in a small subsample of our cohort; therefore, it is unclear whether these results are reliable and require further corroboration. Another major limitation is that our sample showed a sex imbalance, with an overrepresentation of women. This potential recruitment bias may have influenced the results, and it is therefore important to consider how sex affects the contribution of cardiometabolic and AD blood biomarker factors. Another possible limitation is that the biomarkers for Aβ pathology used in this study were based on blood samples. Blood biomarkers appear to be strongly associated with CSF biomarkers [60]; however, they still need to be completely validated in community-based samples. Finally, we did not study other neuroimaging markers, such as atrophy, in a manner complementary to WMHs volume. This approach has been studied previously, allowing for the multimodal comprehension of these markers in the context of the dementia continuum [61].

## Conclusions

Our findings show that WMH in CC dementia-free individuals is more likely to be associated with diastolic blood pressure and changes in glycaemia than Aβ levels. Our results suggest that in the early stages of cognitive decline, microvascular damage, an increase in vascular peripheral resistance, and/or leakage of the BBB arose because of hypertension and hyperglycaemia, which could underlie the origin of WMHs at this step of the dementia continuum. Overall, we suggest adopting a more integrated approach to this vascular phenomenon to achieve a better understanding of these processes. Clinical scenarios, such as metabolic syndrome, or a deeper study of the effects of chronic exposure to glucocorticoids in the brain could be good models for further analysis.

## Supporting information

Online Resources

## Author Contributions

Conceptualization: Patricio Riquelme-Contreras, Michael Hornberger; Methodology: Patricio Riquelme-Contreres, Michael Hornberger, Mizanur Khondoker, Fernando Henriquez, Cecilia Gonzalez-Campo, Florencia Altschuler, Matias Fraile-Vazquez; Formal analysis and investigation: Patricio Riquelme-Contreras, Minzanur Khondoker, Fernando Henriquez, Cecilia Gonzalez-Campo, Florencia Altschuler, Matias Fraile-Vazquez, Writing – original draft preparation: Patricio Riquelme-Contreras, Fernando Henriquez, Cecilia Gonzalez-Campo; Writing – review and editing: Michael Hornberger, Andrea Slachevsky, Pamela Lukasewicz-Ferreira, Bruna Bellaver, Thomas Karikari, Tharick A. Pascoal, Graciela Muniz-Terrera, Patricia Lillo, Funding acquisition: Andrea Slachevsky, Cecilia Okuma, Christian Gonzalez-Billault, Felipe Court, Mauricio Cerda, Patricia Lillo; Supervision: Michael Hornberger and Andrea Slachevsky.

## Ethics Approval

The GERO Chile project was approved by the Scientific Ethics Committee of the Eastern Metropolitan Health Service of Santiago de Chile.

## Informed Consent

Informed consent was obtained from all the participants, as approved by the same committee.

## Consent to publish

Patients signed informed consent regarding publishing their data and photographs.

